# Machine learning modeling of protein-intrinsic features predicts tractability of targeted protein degradation

**DOI:** 10.1101/2021.09.27.462040

**Authors:** Wubing Zhang, Shourya S. Roy Burman, Jiaye Chen, Katherine A. Donovan, Yang Cao, Boning Zhang, Zexian Zeng, Yi Zhang, Dian Li, Eric S. Fischer, Collin Tokheim, X. Shirley Liu

## Abstract

Targeted protein degradation (TPD) has rapidly emerged as a therapeutic modality to eliminate previously undruggable proteins by repurposing the cell’s endogenous protein degradation machinery. However, the susceptibility of proteins for targeting by TPD approaches, termed “degradability”, is largely unknown. Recent systematic studies to map the degradable kinome have shown differences in degradation between kinases with similar drug-target engagement, suggesting yet unknown factors influencing degradability. We therefore developed a machine learning model, MAPD (Model-based Analysis of Protein Degradability), to predict degradability from protein features that encompass post-translational modifications, protein stability, protein expression and protein-protein interactions. MAPD shows accurate performance in predicting kinases that are degradable by TPD compounds (auPRC=0.759) and is likely generalizable to independent non-kinase proteins. We found five features with statistical significance to achieve optimal prediction, with ubiquitination potential being the most predictive. By structural modeling, we found that E2-accessible ubiquitination sites, but not lysine residues in general, are particularly associated with kinase degradability. Finally, we extended MAPD predictions to the entire proteome to find 964 disease-causing proteins, including 278 cancer genes, that may be tractable to TPD drug development.

## Introduction

The most prevalent pathway for selective protein degradation in eukaryotic cells is the Ubiquitin-Proteasome System (UPS), which degrades proteins that are covalently modified with ubiquitin^1–3^. Ubiquitination is orchestrated in three steps by three enzymes. First, ubiquitin is activated by covalent attachment to the active site of an E1 ubiquitin-activating enzyme. Second, the activated ubiquitin is transferred from the E1 enzyme to an E2 ubiquitin-conjugating enzyme. Finally, the proximity induced by an E3 ubiquitin ligase selectively binding to a substrate allows for the covalent transfer of ubiquitin from the E2 enzyme to a lysine residue on the substrate. After repeated rounds of this process, a poly-ubiquitin chain can be formed, which often directs the substrate for degradation by the 26S proteasome^4^.

Targeted protein degradation (TPD) is a novel pharmacologic modality that selectively induces degradation of a protein-of-interest (POI) by chemically repurposing the UPS^5–7^. The TPD molecules (degraders), epitomized by the molecular glues^8, 9^ and PROteolysis TArgeting Chimeras (PROTACs)^5, 10–13^, typically induce the *de novo* ternary complex formation between an E3 ligase and a POI, leading to the ubiquitin transfer to available lysines and subsequent degradation of the POI^14–16^. Unlike traditional inhibitors that target the catalytic binding site on a POI, degraders can induce protein degradation by binding to non-catalytic sites^11, 17, 18^. Therefore, previously undruggable proteins, such as transcription factors (TF), can be targeted by degraders^19, 20^. For example, the FDA-approved immunomodulatory drugs (IMiDs) thalidomide, pomalidomide, and lenalidomide^21–28^ induce degradation of transcription factors IKZF1 and IKZF3 by recruiting them to CRBN^25, 26, 29–32^, the substrate recognition subunit of the E3 ubiquitin ligase complex CUL4-RBX1-DDB-CRBN^33^. Over the last two decades, the TPD field has grown dramatically, with thousands of publicly available degraders developed for over 100 human protein targets^34, 35^. Notably, degraders targeting androgen receptor^36, 37^, oestrogen receptor^38–41^, BCL-XL^42, 43^, Ikaros/Aiolos (IKZF1/3)^44–47^, Helios (IKZF2)^44–46^, and GSPT1^48^ have entered into clinical trials, and degraders targeting STAT3, BRD9, BTK, or TRK will also be tested in patients soon^49^. Despite these advances, it remains challenging to predict which proteins are susceptible and which may be resistant to the TPD approaches.

Chemoproteomic profiling approaches have emerged as a systematic approach to survey protein degradability^50^. Rather than profiling expression of a single protein in response to a selective degrader, these approaches use mass spectrometry to assess the proteome-wide response to treatment with pan-targeting degraders^51–54^. For example, our recent study profiled 91 multi-kinase degraders to assess the degradability of more than 400 protein kinases, identifying more than 200 kinases as degradable^51^. Using a library of pan-HDAC degraders, Xiong *et al*. investigated the degradability of zinc-dependent HDACs^54^. Together these broad-targeted degrader profiling experiments have greatly expanded the known degradable proteome. Unfortunately, chemoproteomic approaches to map degradability are inapplicable for most proteins due to the absence of ligands required for target recruitment to the ligase machinery. Thus, computational prediction of protein degradability offers a potentially practical alternative.

It is widely believed that stable ternary complexes are associated with effective and selective target degradation^15, 16, 53, 55^. A series of computational methods have been introduced to model PROTAC-mediated ternary complex formation^56–59^, which have facilitated the rational and efficient optimization of PROTACs^16, 60^. However, several studies have shown that although some level of binary target engagement and ternary complex formation are necessary for target recruitment and ubiquitin transfer, they are not always sufficient for targeted protein degradation^51–53, 61^. We propose that rather than drug-target interactions driving degradability, features intrinsic to the protein targets could also heavily influence degradability of specific targets. For instance, while ubiquitination is the initiation signal for proteasomal degradation^62–65^, the association between protein degradability and known or potential ubiquitination (Ub) sites in the target protein is poorly understood.

In this study, we developed a machine learning model, MAPD (Model-based Analysis of Protein Degradability), to predict degradability from protein-intrinsic features (Fig. 1). MAPD shows promising performance in predicting degradable kinases by multi-kinase degraders and previously reported targets of PROTAC compounds. We found that a protein’s endogenous ubiquitination potential contributes the most to the degradability predictions. Structural analysis via protein-protein docking revealed the particular importance of E2-accessible Ub sites in determining degradability. Using MAPD, we have expanded our predictions to the human proteome to map protein tractability to TPD approaches. Our results are available at http://mapd.cistrome.org/, which could be a valuable resource for guiding target prioritization towards tractable TPD targets.

**Fig. 1.**
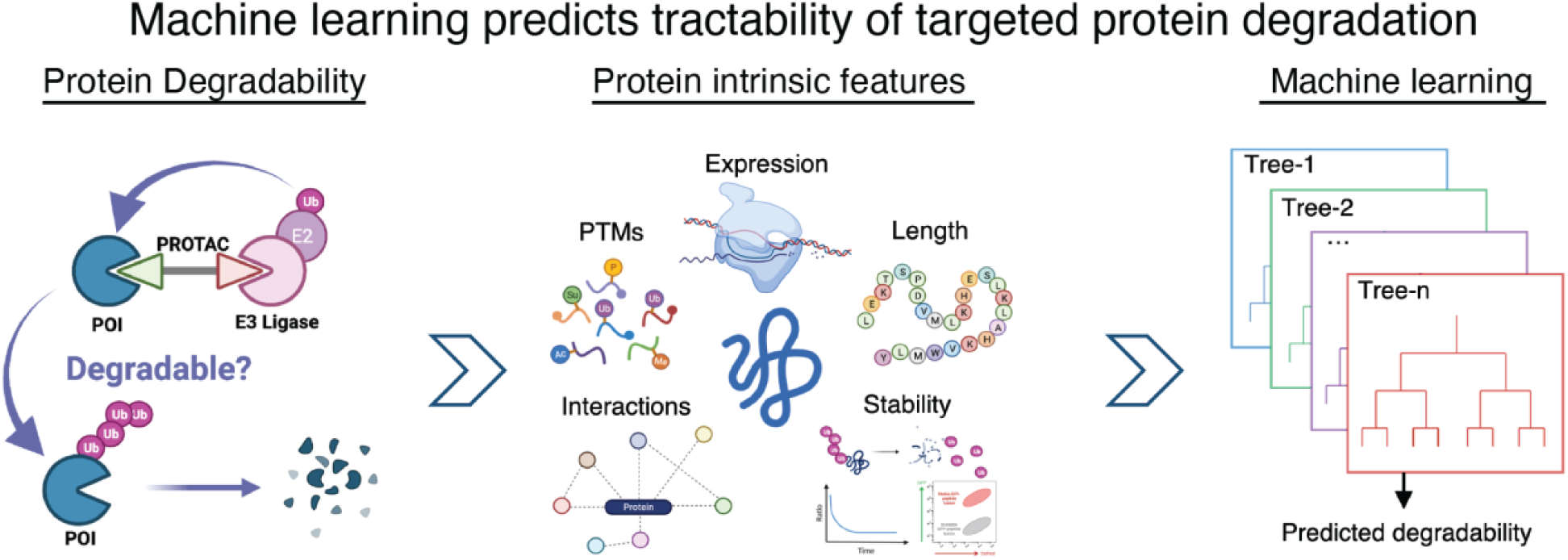
Study overview. The ubiquitin-proteasome system can be repurposed by a PROTAC (Proteolysis Targeting Chimera) or other small molecule to degrade a protein of interest (POI). However, it remains to be answered which proteins are amenable to this approach (left). Here, we associated kinase degradability with protein-intrinsic features spanning protein expression, post-translational modifications, protein length, protein-protein interactions, protein stability, and protein half-life to identify predictive factors (middle). Based on the predictive features, we developed a machine learning model to predict protein degradability (right).

## Results

### Kinase degradability is associated with features intrinsic to the target

Substantial efforts have been invested in the optimization of degraders for any particular target with no guarantee that a successful compound will be found^66, 67^. Our previous chemoproteomic study of the protein kinome indicates that drug-target engagement is insufficient to predict which kinases can be degraded^51^, suggesting unexplained factors influencing protein degradability. In this study, we explored factors intrinsic to POIs that may influence their degradability by comparing kinases that all have drug-target engagement, but differ in multi-kinase degrader-induced degradation. We first selected highly- and lowly-degradable kinases based on the number of multi-kinase degraders found to degrade each POI (Fig. 2a), with an additional requirement of high frequency of detection in the underlying global proteomic experiments (Extended Data Fig. 1a). We next collected protein features that may be predictive of kinase degradability, including post-translational modifications (PTMs), protein stability, protein-protein interaction (PPI), protein expression, etc. (Supplementary Table 1). Often features within a category are highly correlated with each other, while features between categories tend to provide independent information (Fig. 2b).

**Fig. 2.**
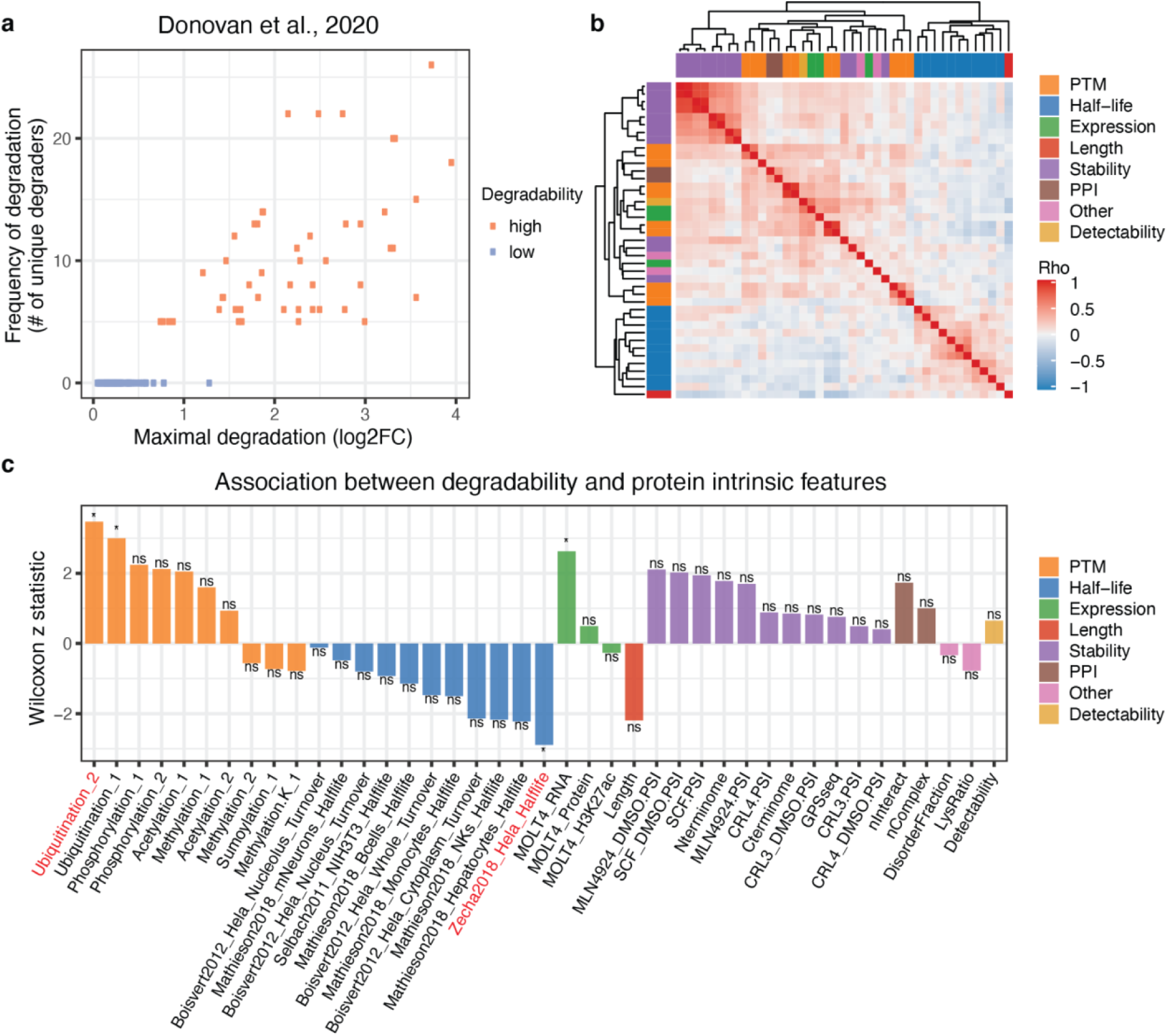
Kinase degradability is associated with features intrinsic to the target. **a**, Dot plot showing the frequency of degradation and maximal degradation of protein kinases induced by multi-kinase degraders from the Donovan et al. study. Orange dots represent the kinases with high degradability, and light blue dots represent the kinases with low degradability. **b**, Pairwise Spearman’s correlation of 42 protein-intrinsic features spanning protein stability, post-translational modification (PTM), protein-protein interaction (PPI), protein length, protein half-life, protein expression, protein detectability and others. **c**, Bar diagram showing the association between degradability of kinases and their features. The x-axis shows the abbreviated name of protein-intrinsic features (see Supplementary Table 1 for full details), and the y-axis shows the Wilcoxon z-statistics indicating the association between protein degradability and each protein-intrinsic feature (*=FDR<0.05).

To identify features associated with protein degradability, we compared highly- and lowly- degradable kinases using a Wilcoxon rank-sum test. Compared to lowly-degradable kinases, the highly-degradable kinases have a significantly higher proportion of lysine residues that have reported ubiquitination events from the PhosphoSitePlus database^68^ (hereafter referred to as ubiquitination potential) (p=5.2e-4; Fig. 2c, S1b-c). The ubiquitination potential likely reflects a protein’s endogenous capacity to be ubiquitinated since the ubiquitination events are from cell lines in the absence of degrader treatment^69^. Notably, the percentage of lysine residues on POIs are not significantly different (Extended Data Fig. 1d). Besides ubiquitination potential, mRNA expression of a POI in the assayed cell lines is positively associated with protein degradability (Fig. 2c, S1e), suggesting that profiling in more cell contexts might be advantageous. Furthermore, we observed an enrichment of proteins with lower half-life in the highly-degradable group (Fig. 2c, S1f). Given that protein half-life was not correlated with ubiquitination potential (Extended Data Fig. 1g), this indicates an independent signal for predicting protein degradability. Collectively, these results suggest that features intrinsic to protein targets might influence their degradability.

### Development of Model-based Analysis of Protein Degradability (MAPD)

We next sought to build a machine learning model, named Model-based Analysis of Protein Degradability (MAPD), to combine multiple features associated with protein degradability into a single score. Towards this end, we tested six commonly used machine learning methods, including naive bayes (NB), k-nearest neighbor (KNN), logistic regression, linear-kernel support vector machine (svmLinear), radial kernel support vector machine (svmRadial), and random forest (RF). Because of the redundancy of protein-intrinsic features, we performed forward feature selection for each method (Methods), which iteratively selects the best-performing features (Supplementary Table 2) until the model performance plateaus^70^. By evaluating performance using cross-validation, the RF model outperformed other models with an area under the Precision-Recall Curve (auPRC) of 0.759 (Fig. 3a) and area under the receiver operating characteristic curves (auROC) of 0.773 (Extended Data Fig. 2a). Therefore, all further analyses are based on the RF model implementation.

**Fig. 3.**
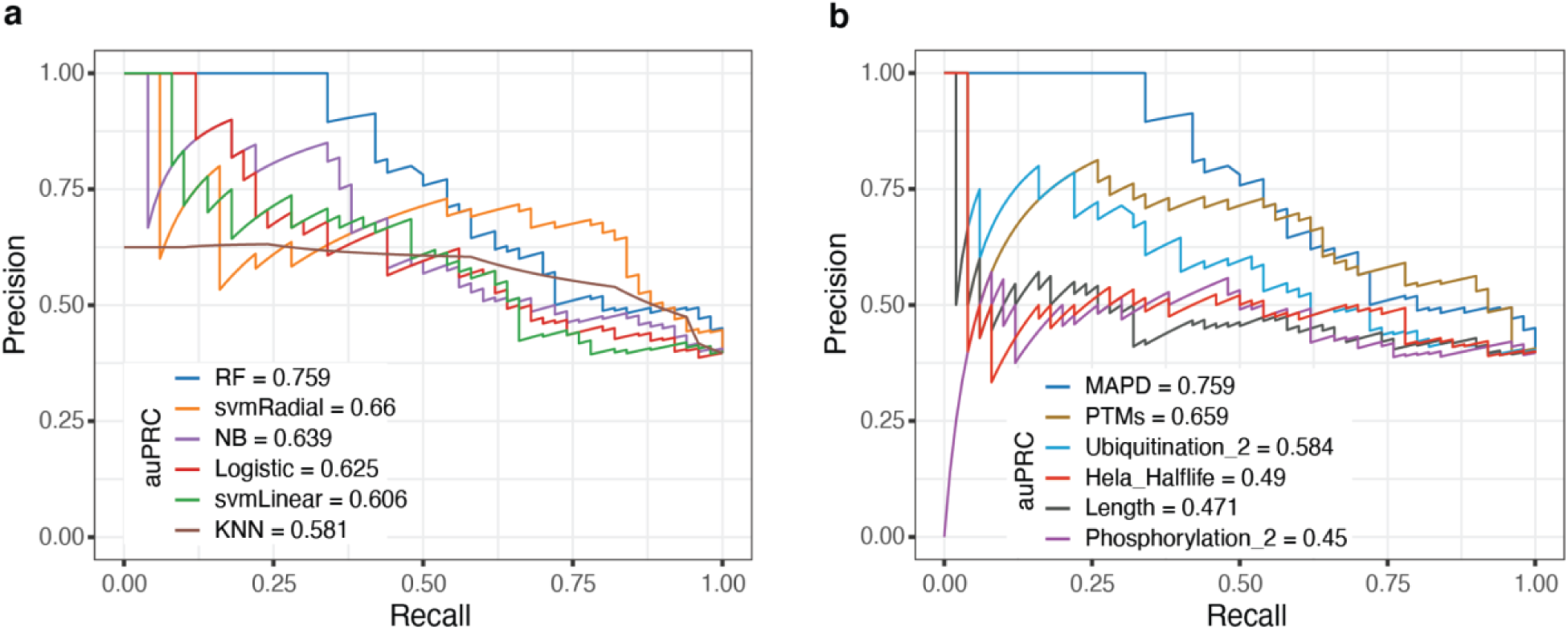
Development of Model-based Analysis of Protein Degradability (MAPD). **a**, Precision-Recall curves that show the performance of six machine learning models based on 20-fold cross-validation. RF indicates the random forest model, svmRadial indicates the radial-kernel support vector machine model, NB indicates the naive bayes model, Logistic indicates the logistic regression model, svmLinear indicates the linear kernel support vector machine model, and KNN indicates the k-nearest neighbor model. **b**, Precision-Recall curves that show the performance of MAPD and models trained on individual features or combination of features. ‘PTMs’ indicates the model trained on the combination of ubiquitination potential (Ubiquitination_2), acetylation potential (Acetylation_1), and phosphorylation potential (Phosphorylation_2). ‘Ubiquitination_2’ indicates the model trained on ubiquitination potential. ‘Hela_Halflife’ indicates the model trained on a single feature describing half-life in Hela cells from Zecha *et al*. ‘Length’ indicates the model trained on protein length. ‘Phosphorylation_2’ indicates the model trained on phosphorylation potential.

Five protein-intrinsic features were identified as important in the MAPD model, including ubiquitination potential, phosphorylation potential, protein half-life, acetylation potential, and protein length (Extended Data Fig. 2b), in order of importance. Next, we compared the performance of MAPD to models that were trained on each individual feature using cross-validation. Consistent with the highest importance of ubiquitination potential in MAPD, the model trained on the ubiquitination potential showed the highest auPRC (0.584) and auROC (0.663) among all other single-featured models (Fig. 3b, Extended Data Fig. 2c). Interestingly, the combination of the three PTM features (ubiquitination, phosphorylation, and acetylation) seem to achieve higher auPRC (0.659) and auROC (0.753) than ubiquitination potential alone (p= 0.058, Delong’s test) (Fig. 3b, Extended Data Fig. 2c). This suggests that the general propensity of a protein to be post-translationally modified might be predictive of protein degradability.

### MAPD shows good performance in predicting kinase degradability

To evaluate the robustness of MAPD, we assessed the degradability of the kinome, with the predictions for training kinases collected from the 20-fold cross-validation to avoid inflating the performance assessment. We first examined the degradability of kinases profiled in Donovan *et al*.^51^ and found significantly higher MAPD scores of degradable kinases than other kinases engaged by multi-kinase degraders (Extended Data Fig. 3a). This trend is also consistent for specific degraders, such as TL12-186 and SK-3-91 (Extended Data Fig. 3a), although with less significance due to the smaller number of POIs in these datasets. Based on a threshold with the best cross-validation accuracy, MAPD identified 382 highly-degradable kinase/kinase-related proteins, covering 78.8% (171/217) experimentally degradable kinases^51^ (Fig. 4a). Consistent with the low MAPD scores, the remaining 21.2% kinases have a low frequency of degradation (Extended Data Fig. 3b). Furthermore, within all experimentally degraded kinases, MAPD scores show considerable correlation with their frequency of degradation by multi-kinase degraders (p=5.51e-6) (Fig. 4b), indicating the capability of MAPD in prioritizing highly-degradable targets. We next examined the overlap of degradable targets from MAPD and curated protein targets with reported PROTACs in databases (PROTAC-DB^34^ and PROTACpedia^35^). Although some PROTAC targets were missed (Supplementary Table 3), MAPD successfully identified 77% (50/65) of kinase targets (Fig. 4a), supporting its ability in distinguishing degradable kinases from other kinases. In addition, MAPD recovered 14 PROTAC targets that were not identified by Donovan *et al.*^51^ (Fig. 4a), which highlights how computational methods can be complementary to high-throughput experimental approaches.

**Fig. 4.**
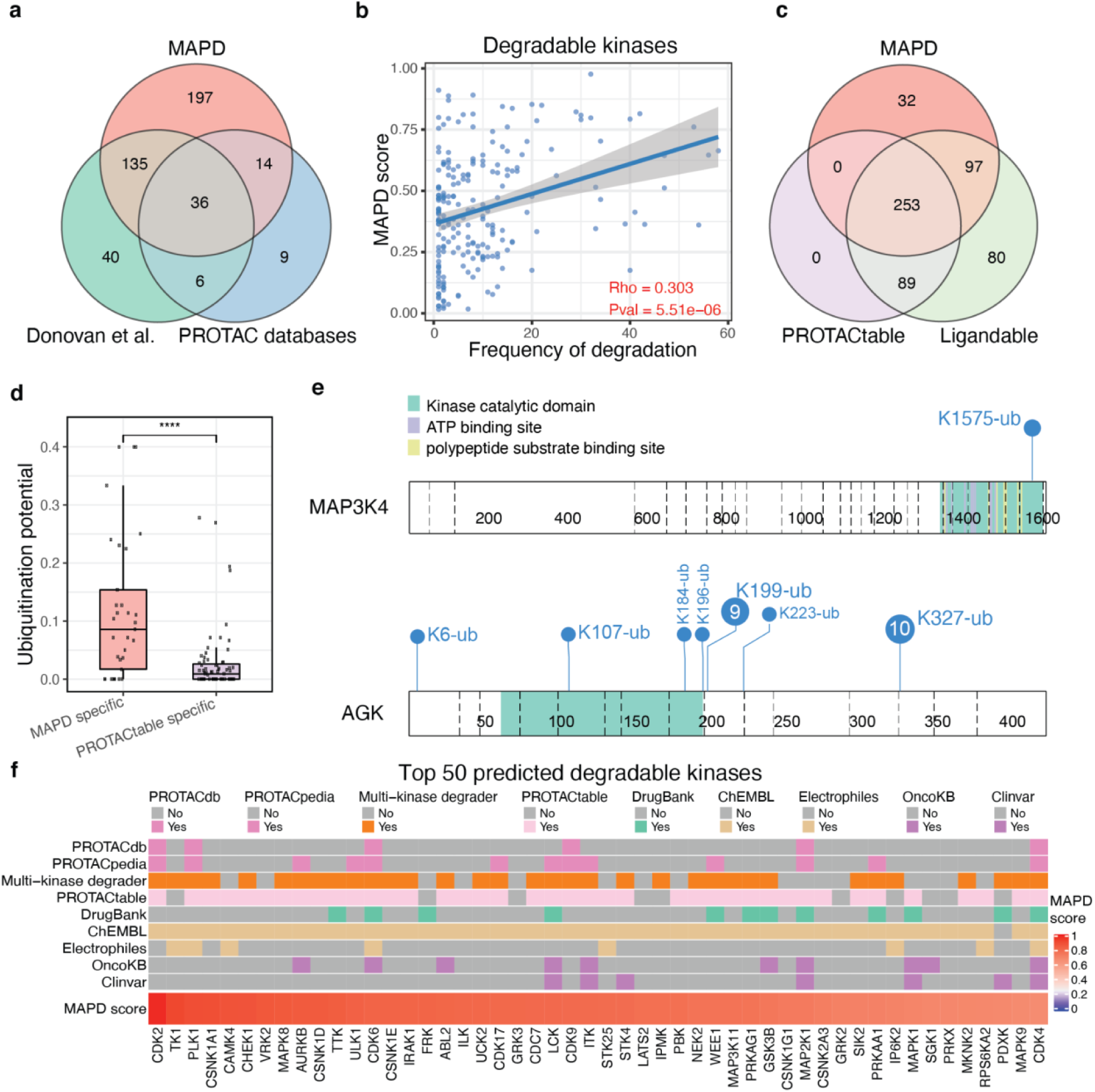
MAPD shows good performance in predicting kinase degradability. **a**, Venn diagram showing the overlap between kinases degraded by multi-kinase degraders from Donovan *et al*., PROTAC targets reported in PROTAC databases (including PROTAC-DB and PROTACpedia), and degradable kinases identified by MAPD. **b**, Scatter plot showing the Spearman correlation between MAPD scores and frequency degradation of all degradable kinases from Donovan *et al*. **c**, Venn diagram showing the overlap between degradable kinases identified by MAPD, PROTACtable kinases, and ligandable kinases. **d**, Box plot showing ubiquitination potential (proportion of lysine residues with reported ubiquitination events in the PhosphoSitePlus) of MAPD-specific targets and PROTACtable-specific targets. **e**, Lollipop diagram showing the reported Ub sites in MAP3K4 (PROTACtable-specific target) and AGK (MAPD-specific target). The number in the circles indicates the number of references for each Ub site in PhosphoSitePlus and the blank circle indicates that only one reference is available. The blue text near the circle indicates the location of the Ub site. **f**, Heatmap showing annotations of the top 50 predicted degradable kinases, with MAPD scores shown at the bottom. ‘PROTACdb’ and ‘PROTACpedia’ indicate whether a kinase has a developed degrader reported in the respective databases. The ‘Multi-kinase degrader’ indicates whether a protein is degraded by the multi-kinase degrader. ‘DrugBank’ indicates whether a protein has FDA approved drug recorded in the DrugBank database. ‘ChEMBL’ indicates whether a protein has ligands recorded in the ChEMBL database. ‘Electrophiles’ indicate whether a protein has ligandable cysteines from the SLCABPP (Streamlined Cysteine Activity-Based Protein Profiling). The ‘OncoKB’ indicates whether a protein is considered as a cancer gene in the OncoKB database. The ‘ClinVar’ indicates whether the protein is associated with a disease in the ClinVar database (****=p<0.0001).

A binder of the target protein is required in the design of TPD molecules, so the propensity of a POI to be bound by a small molecule, also called ligandability, is relevant to tractability of the POI by TPD molecules. Here, we leveraged knowledge of existing small molecules to refine MAPD predictions. A protein is considered ligandable if it has at least one ligand reported in PROTAC-DB^34^, PROTACpedia^35^, DrugBank^71^, ChEMBL^72^ or SLCABPP (Ligandable Cysteine Database)^73^ (Extended Data Fig. 3d). Out of the 519 ligandable kinases, MAPD identified 350 degradable kinases, including 74% (253/342) PROTACtable targets and 97 targets specifically identified by MAPD (Fig. 4c). PROTACtable was introduced in a recent perspective article^74^ that qualitatively assigned tractable TPD targets based on ligand records in ChEMBL and a rule-based approach that only considers whether certain protein annotations are available. We observed a significantly lower ubiquitination potential of PROTACtable-specific targets than MAPD-specific targets (Fig. 4d). For example, MAP3K4, a PROTACtable-specific target, has only one reported Ub site despite being a particularly long protein with 103 lysines^68^ (Fig. 4e). In contrast, the MAPD-specific target, AGK, is extensively ubiquitinated despite its short length (Fig. 4e). Experimental data showed that AGK was degraded sufficiently by multi-kinase degraders^51^ while MAP3K4 was not despite its strong target engagement by a multi-kinase degrader^52^. These examples highlight a potential advantage of MAPD by quantitatively assessing protein degradability.

In total, MAPD identified 132 disease-relevant kinase targets, including 72 cancer genes in OncoKB and 60 kinases associated with other diseases reported in the ClinVar database ^75, 76^ (Extended Data Fig. 3e). These kinases could be prospective targets for development of degraders (Supplementary Table 3). The most degradable kinases include targets with developed PROTACs^34, 35^, such as CDK2, PLK1, CDK6, CDK9 and CDK4, and other promising targets, such as TK1, CSNK1A1, CHEK1, MAPK8, and AURKB that are degraded by multi-kinase degraders^51, 52^ (Fig. 4f).

### MAPD predicts proteome-wide degradability

We hypothesized that MAPD might also predict the degradability of non-kinase proteins. To test this, we collected 65 non-kinase targets with publicly available degraders reported in PROTAC databases^34, 35^. These PROTAC targets had significantly higher MAPD scores than other drug targets from DrugBank^71^ (Fig. 5a). To further corroborate this finding, we collected a list of TFs, such as Ikaros (IKZF1) and Aiolos (IKZF3), that are frequently degraded by thalidomide analog (IMiD)-based degraders^32, 51^. The MAPD scores of these TFs showed significant correlation with their observed frequency of degradation (p=0.022) (Fig. 5b). Additional TFs have also been targeted by TPD molecules^20, 77, 78^, such as degraders for AR^38, 79–81^ and ER^82–86^ that have entered into clinical trials. With the exception of BCL6 which has few reported Ub sites, MAPD correctly predicts the high degradability of most TF PROTAC targets (Fig. 5c). Taken together, these results indicate that MAPD is generalizable to POIs outside of the kinome.

**Fig. 5.**
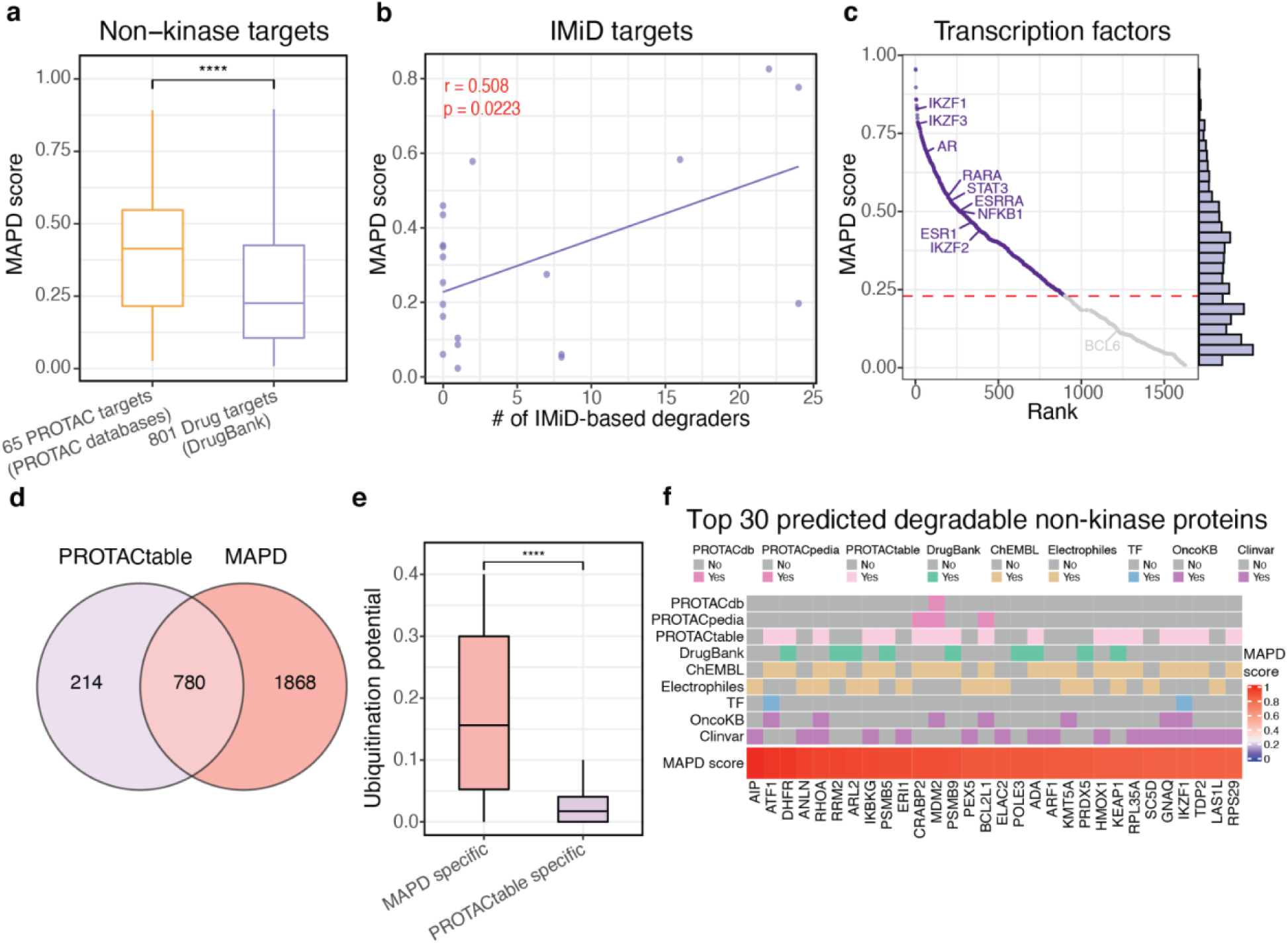
MAPD predicts degradability proteome-wide. **a**, Box plot showing the MAPD scores of non-kinase PROTAC targets from PROTAC databases (including PROTAC-DB and PROTACpedia) and other non-kinase drug targets from DrugBank. **b**, Scatter plot showing the MAPD scores and the frequency of degradation of IMiD targets by CRBN-recruiting degraders from Donovan *et al*. **c**, Ranked dot plot showing the MAPD scores of human transcriptional factors (TF). TFs with reported degraders are labeled on the figure. The histogram at right shows the distribution of MAPD scores of all human TFs and the red dashed line shows the threshold for identifying degradable proteins by MAPD. **d**, Venn diagram showing the overlap of degradable non-kinase proteins between MAPD predictions and PROTACtable genome. **e**, Box plot showing the ubiquitination potential (proportion of lysines with reported ubiquitination events in the PhosphoSitePlus) in MAPD-specific targets and PROTACtable genome-specific targets. **f**, Heatmap showing annotations of the top 30 predicted degradable non-kinase proteins, with MAPD scores shown at the bottom. ‘PROTACdb’ and ‘PROTACpedia’ annotations indicate whether a kinase has a developed degrader reported in the respective databases. The ‘Multi-kinase degrader’ indicates whether a protein is degraded by the multi-kinase degrader. ‘DrugBank’ indicates whether a protein has FDA approved drug recorded in the DrugBank database. ‘ChEMBL’ indicates whether a protein has ligands recorded in the ChEMBL database. ‘Electrophiles’ indicate whether a protein has ligandable cysteines from the SLCABPP (Streamlined Cysteine Activity-Based Protein Profiling). ‘OncoKB’ indicates whether a protein is considered as a cancer gene in the OncoKB database. ‘ClinVar’ indicates whether the protein is associated with a disease in ClinVar database (****=p<0.0001).

Given the robust performance of MAPD, we next applied MAPD proteome-wide to systematically score all proteins outside of the kinome. MAPD predicted 2,648 degradable targets out of 4,137 ligandable non-kinase proteins (Extended Data Fig. 4a,b), which was two-fold more than PROTACtable^74^ (Fig. 5d). The MAPD-specific targets again had significantly higher levels of ubiquitination potential than the PROTACtable-specific targets (Fig. 4e). We further identified 832 disease-relevant non-kinase targets that are amenable to TPD (Extended Data Fig. 4c and Supplementary Table 4). Of these, 206 proteins are considered as oncogenic genes by OncoKB and 626 proteins are associated with other human diseases reported in the ClinVar database^75, 76^ (Extended Data Fig. 4c). The top predicted degradable targets include known PROTAC targets, such as MDM2 and BCL-XL (BCL2L1), and other potentially degradable targets. DHFR, one of the top-ranking targets, has been successfully degraded by a hydrophobic tagging probe consisting of a hydrophobic moiety Boc3Arg and a DHFR non-covalent binding ligand TMP^87^. RHOA, RHOB, and RHOC are also predicted to be degradable, which have been previously reported to be degraded by F-box-intracellular single-domain antibodies^88^. These results suggest potential opportunity for future TPD efforts (Fig. 5f).

### The E2-accessibility of Ub sites is associated with protein degradability

Given that ubiquitination potential was the most important feature in MAPD, we hypothesized that structural properties of Ub sites could be informative of protein degradability. To test this hypothesis, we first grouped Ub sites according to their structural properties (Supplementary Table 4) such as secondary structure, relative solvent accessibility, or flexibility (as defined by B-factor)^89^. We then examined the association between protein degradability and the number of Ub sites in each group using a Wilcoxon z-statistic. Among annotated secondary structures, the number of Ub sites in loop regions showed modestly higher association with protein degradability relative to the total number of Ub sites (Extended Data Fig. 5a). However, neither relative solvent accessibility nor flexibility of Ub sites improved the association with protein degradability (Extended Data Fig. 5b,c). These data suggest that local structural properties of a Ub site provide limited information for predicting protein degradability.

We next investigated the property of Ub sites that facilitates the transfer of ubiquitin from the attached E2 enzyme to the POI in degrader-mediated ternary complexes. We reasoned that quantifying the accessibility of Ub sites to the E2 enzyme might be predictive of protein degradability. As most degraders in the chemoproteomics study were based on the CRBN substrate receptor, we examined this hypothesis by computationally docking 251 target kinases with experimental structures onto CRBN–IMiD (Extended Data Fig. 6a). We examined the 200 top-scoring structural models for each POI and removed those where it was not feasible to fit a PROTAC (Extended Data Fig. 6b). Due to the high flexibility of the CUL4 arm, the attached E2 can transfer ubiquitin to any site in a broad ubiquitination zone^90^, hence all Ub sites in the spatial quadrant facing the E2 were considered accessible to the E2 (Fig. 6a, Extended Data Fig. 6c). We then defined E2 accessibility as the fraction of top-scoring models in which the Ub site was accessible to the E2 enzyme (Fig. 6a, Extended Data Fig. 6c, Supplementary Table 4). In comparison to the total number of Ub sites in the structure of the POI, the E2-accessible Ub sites showed a more significant positive association with protein degradability (Fig. 6b, Extended Data Fig. 7a). In contrast, the number of E2-accessible lysine residues on the POIs does not show significant association with their degradability (Extended Data Fig. 7a,b). Together, these results suggest that lysines with detected ubiquitination events are more amenable to TPD. To further assess whether E2-accessibility was independently useful, we randomly shuffled reported Ub sites among all available lysine residues within a protein. Consistent with our initial finding, E2-accessible Ub sites were significantly more associated with protein degradability than expected based on the total number of Ub sites in each protein (p=0.0064; Fig. 6c).

**Fig. 6.**
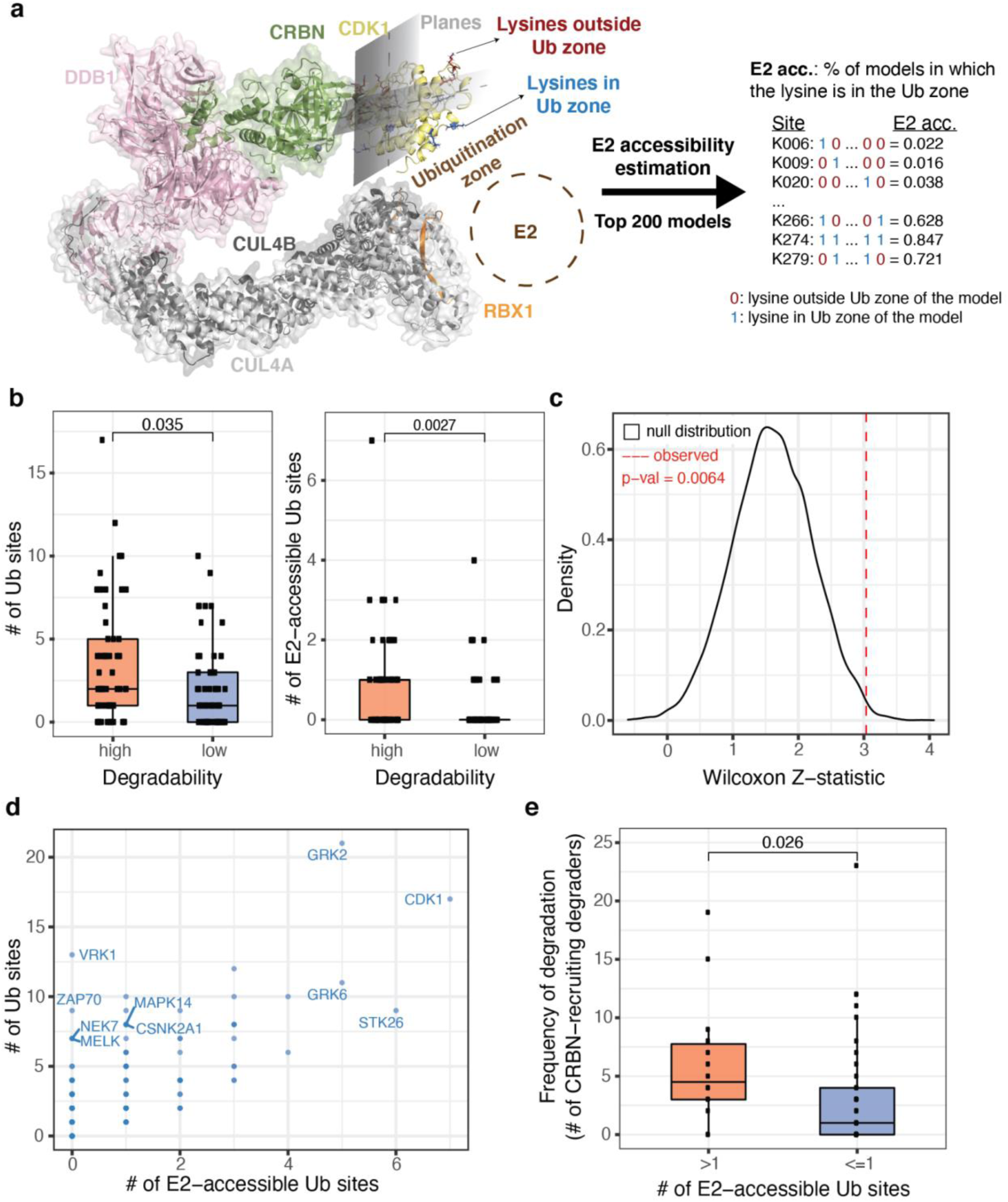
E2-accessibility of Ub sites is associated with protein degradability. **a**, Diagram showing how to estimate accessibility of lysine/Ub sites to E2 enzyme in degrader-induced ternary complex. The model of CDK1 (4Y72) was docked to the CRBN-Lenalidomide structure (PDB: 5FQD), which is shown as an example. The E3 ubiquitin ligase complex consists of CRBN, DDB1, CUL4A, and CUL4B, shown in green, pink, light gray and gray, respectively. The CDK1 is the target protein, shown in yellow. The RBX1 fragment (shown in orange) was used to estimate the position of the E2 enzyme and corresponding ubiquitination zone in the target protein. Lysine/Ub sites in the ubiquitination zone were estimated by drawing two planes with respect to the position of CRBN and the target kinase. The sites lying in the quadrant facing the putative position of the E2, estimated by the placement of RBX1 are considered accessible. The predicted E2-accessible and E2-inaccessible lysine residues are highlighted in blue and red, respectively. For each target protein, 200 top-scoring feasible models are selected for evaluating the accessibility of lysine residues to E2 enzyme. For each Ub site, the fraction of feasible models with the site in the ubiquitination zone was estimated as its E2 accessibility. **b**, Box plot showing the association of kinase degradability with total number of Ub sites (left) and E2-accessible Ub sites (right) in the kinases. The E2-accessible Ub sites (E2 accessibility >=0.5) were defined as the Ub sites lying in the ubiquitination zone of more than 50% feasible models. **c**, Density plot showing the null distribution of Wilcoxon z-statistics generated by shuffling Ub sites among all lysine residues for 10,000 times. The red dashed line indicates the observed Wilcoxon z-statistic representing the association between protein degradability and the number of E2-accessible Ub sites (E2 accessibility >=0.5). **d**, Dot plot showing the total number of resolved Ub sites and the number of E2-accessible Ub sites (E2 accessibility >=0.5). **e**, Box plot showing the number CRBN-recruiting degraders that degrade kinases with high (>1) and low (<=1) level of E2-accessible Ub sites. All kinases involved in this analysis have at least two reported Ub sites, which reduces the confounding effect derived from the difference in the total number of Ub sites.

We observed an overall positive correlation between the total number of Ub sites and E2 accessible Ub sites on kinases (Fig. 6d), and noticed some POIs with outlier levels of E2 - accessible and total Ub sites. For example, CDK1 had a high fraction of E2-accessible Ub sites (Fig. 6d, Extended Data Fig. 7c), consistent with its frequent degradation by multi-kinase degraders^51^. Therefore, we hypothesize that similar proteins, such as GRK2, GRK6, and STK26, are promising targets for developing future TPD drugs if they had drug-target engagement (Fig. 6d). In contrast, some kinases, such as VRK1, ZAP70, NEK7, and MAPK14, had a low number of E2-accessible Ub sites, despite having a high number of total Ub sites (Fig. 6d). As expected, these kinases have significantly lower frequency of degradation by CRBN-recruiting multi-kinase as measured by Donovan *et al.*^51^ (Fig. 6e).

Finally, we created an interactive web platform (http://mapd.cistrome.org), which incorporates protein-intrinsic features, MAPD predictions, E2 accessibility of Ub sites in select proteins, ligandability, and disease associations. This platform could enable rational prioritization of degradable targets for developing degraders by the TPD community. Moreover, we implemented MAPD as a R package (https://github.com/liulab-dfci/MAPD), which allows researchers to extend our analysis when more chemoproteomic profiling data and/or protein features are available in the future.

## Discussion

Despite the growth in the number of targeted protein degraders, it remains challenging to predict which proteins are tractable to this approach. In this study, we investigated the degradability of kinases and their correlation with features intrinsic to protein targets. By developing a machine learning model, MAPD (Model-based Analysis of Protein Degradability), we identified five features predictive of kinase degradability, including the ubiquitination potential, acetylation potential, phosphorylation potential, protein half-life and protein length. Systematic benchmarking indicates that MAPD can well predict kinase degradability and is also applicable to proteins outside of the kinome. By integrating MAPD predictions and ligand information of POIs, we prioritized disease-associated degradable proteins as TPD drug targets.

Ternary complex formation is thought to be the most important factor in determining the degradability of protein targets^53, 55–59^. However, our analysis found that protein degradability can also be heavily influenced by protein-intrinsic features, especially the protein’s endogenous ubiquitination potential. By modeling the structural relationship between target proteins and E2 enzyme, we found that protein degradability is highly correlated with the availability of E2 - accessible Ub sites. Thus, checking the protein-intrinsic features, especially the availability of E2- accessible Ub sites, might be crucial for selecting protein targets or E3 recruiters before a TPD drug discovery project.

Our study has several limitations. First, our analysis revealed protein-intrinsic features, such as ubiquitination potential and protein half-life, associated with protein degradability, but it remains to be answered how they influence protein degradability. Second, although our model had the potential to identify degradable non-kinase targets, it showed biased predictions for some proteins (e.g., BRD4, BCL6, HDAC6, and HDAC3) with poorly detected Ub sites or missing feature data. Therefore, a careful consideration of feature data is important when interpreting the prediction results. Lastly, while E2-accessible Ub sites are important in determining protein degradability, we didn’t incorporate this feature into MAPD. One reason is that most proteins don’t have experimentally solved protein structure with known ligandable pockets, which is required for protein docking models. The release of highly accurate predicted protein structures generated with AlphaFold may offer a great opportunity for researchers to address this problem in the future^91^.

Our study also reveals several research directions deserving future study to advance the field. First, computational and experimental studies investigating why certain lysines seem more susceptible to ubiquitination than others could improve the predictions for degradability by MAPD. Second, more extensive proteomic profiling of protein-intrinsic features and induced protein degradation by multi-target degraders in disease-relevant cell lines or tissues could facilitate the understanding of cell-type-specific protein degradability and further accelerate the development of TPD drugs for diseases. Finally, we envision that future computational methods will not only improve the prediction of protein degradability, but also predict the functional consequence of degradation of disease-causing proteins.

## Acknowledgements

This study was supported by grants from the Breast Cancer Research Foundation (BCRF-19-100 to X.S.L.), the Mark Foundation for Cancer Research (Mark Foundation Emerging Leader Award 19-001-ELA to E.S.F.), the NIH (R01CA218278 and R01CA214608 to E.S.F.), and Cancer Research Institute (Irvington Postdoctoral Fellowship CRI 3442 to S.S.R.B.). C.T. is a Damon Runyon Fellow supported by the Damon Runyon Cancer Research Foundation (DRQ-04-20). We acknowledge the Research Computing Group at Harvard Medical School and Dana-Farber Cancer Institute for cluster time, and the SBGrid consortium for structural biology software. We also would like to thank Dr. Chris Sander for helpful suggestions on this study.

## Author Contributions

W.Z., C.T., and X.S.L. conceived of the study. W.Z., S.S.R.B., K.A.D., B.Z., E.S.F., C.T., and X.S.L. drafted and edited the manuscript. W.Z. developed the computational methods. W.Z. and S.S.R.B. performed the protein structural analysis. J.C. developed the interactive website. K.A.D. contributed degradability data. Y.C., Z.Z, Y.Z, and D.L participated in discussions.

## Extended Data Figures

**Extended Data Fig. 1.**
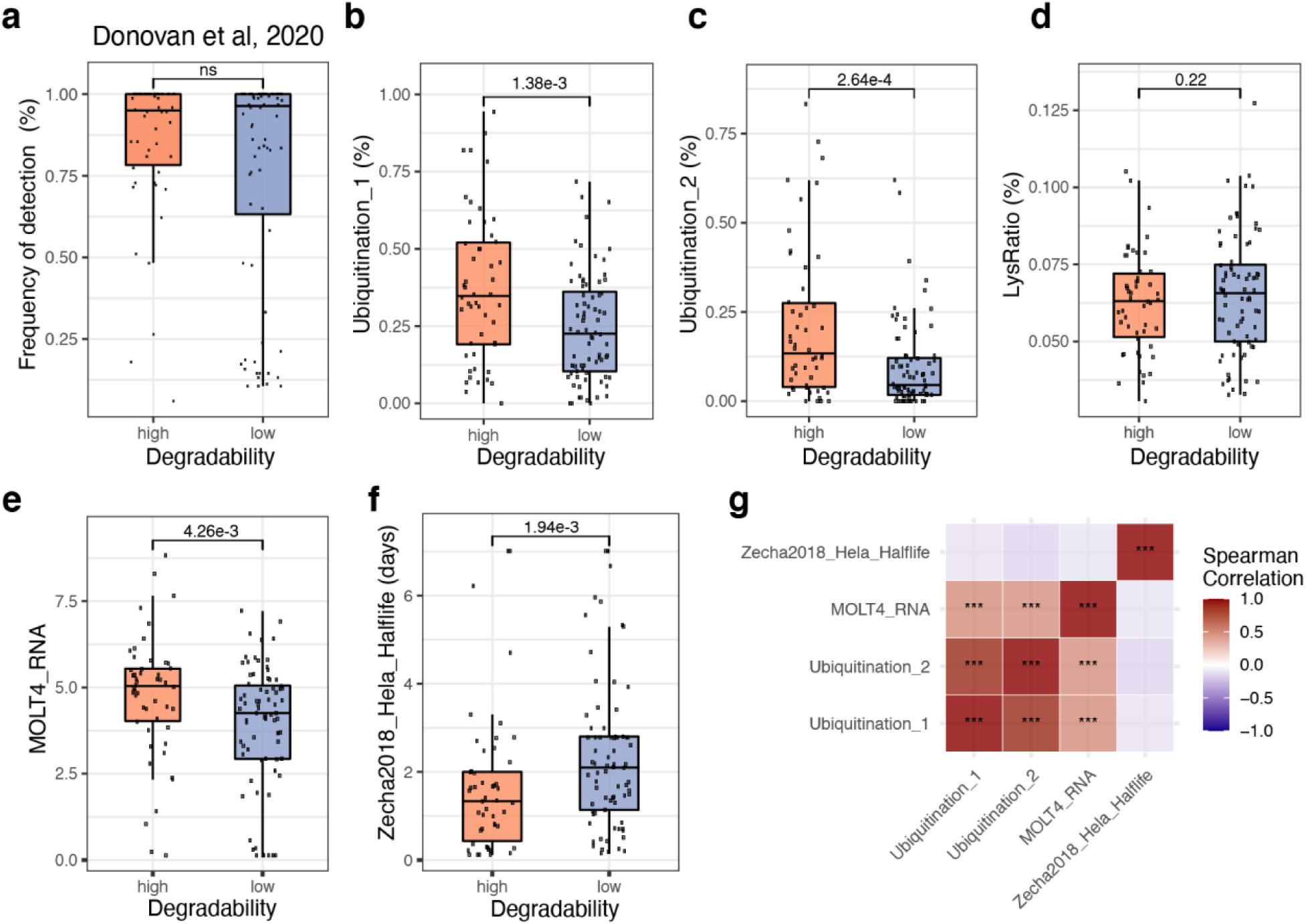
Kinase degradability is associated with features intrinsic to the target. Related to Fig. 2. **a-f**, Box plot showing difference between high-degradability and low-degradability kinases for (**a**) frequency of detection in the chemoproteomic data from Donovan *et al*. study, (**b**) proportion of lysines with at least one reported ubiquitination event in the PhosphoSitePlus, (**c**) proportion of lysines with at least two reported ubiquitination events in the PhosphoSitePlus, (**d**) fraction of lysine residues, (**e**) mRNA expression in the MOLT4 cell line, and (**f**) protein half-life in Hela cells. **g**, Heatmap showing the pairwise Spearman correlation of the four protein-intrinsic features. **h**, Heatmap of Wilcoxon z statistics indicating the association between protein degradability and protein-intrinsic features of kinases in each family. The x-axis shows the abbreviated name of protein-intrinsic features (see Supplementary Table 1 for full details). The y-axis shows the kinase family with the number of highly-degradable (H) and lowly-degradable (L) kinases shown in the label. The color shows the Wilcoxon z-statistics indicating the association between protein degradability and each protein-intrinsic feature (ns=p>0.05, *=p<0.05, **=p<0.01, ***=p<0.001).

**Extended Data Fig. 2.**
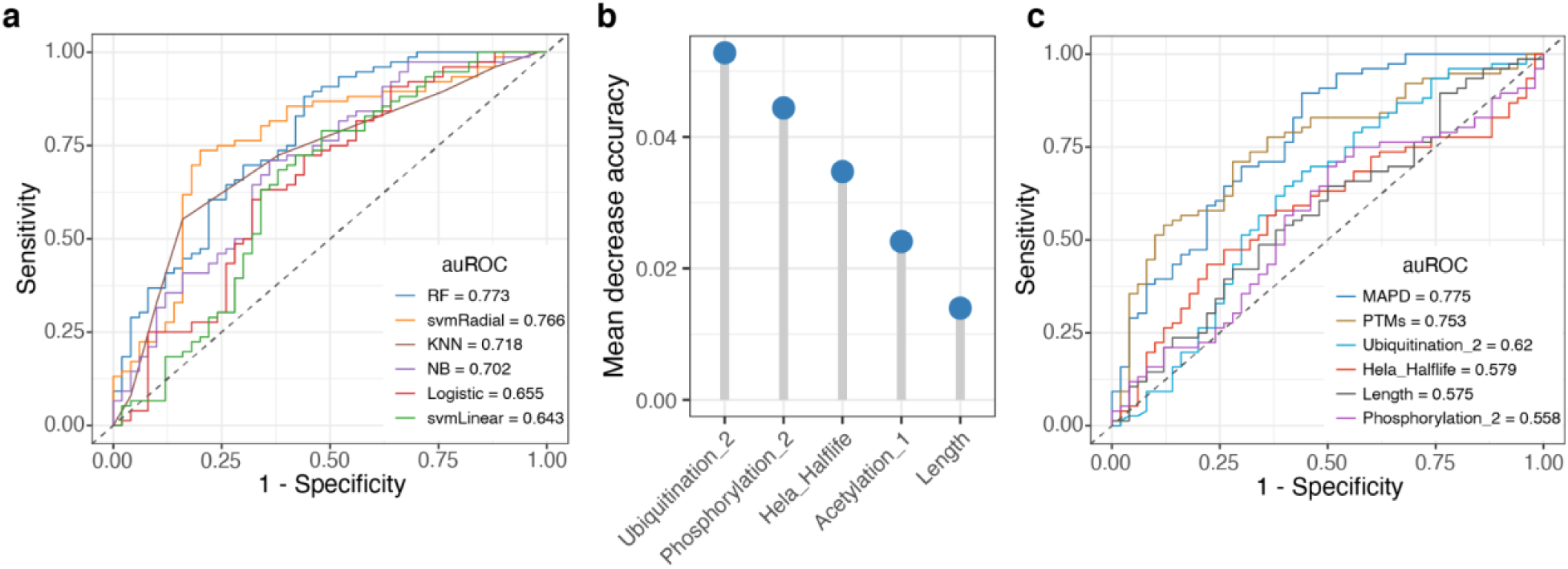
Development of Model-based Analysis of Protein Degradability (MAPD). Related to Fig. 3. **a**, ROC curves (receiver operating characteristics curves) showing the performance of six machine learning models in predicting kinase degradability based on 20-fold cross-validation. **b**, Importance of five features in the MAPD revealed by mean decrease accuracy metric that measures how much accuracy the model losses by excluding each feature from the model. **c**, ROC curves (receiver operating characteristics curves) showing the performance of MAPD and models trained on a subset of features based on 20-fold cross-validation.

**Extended Data Fig. 3.**
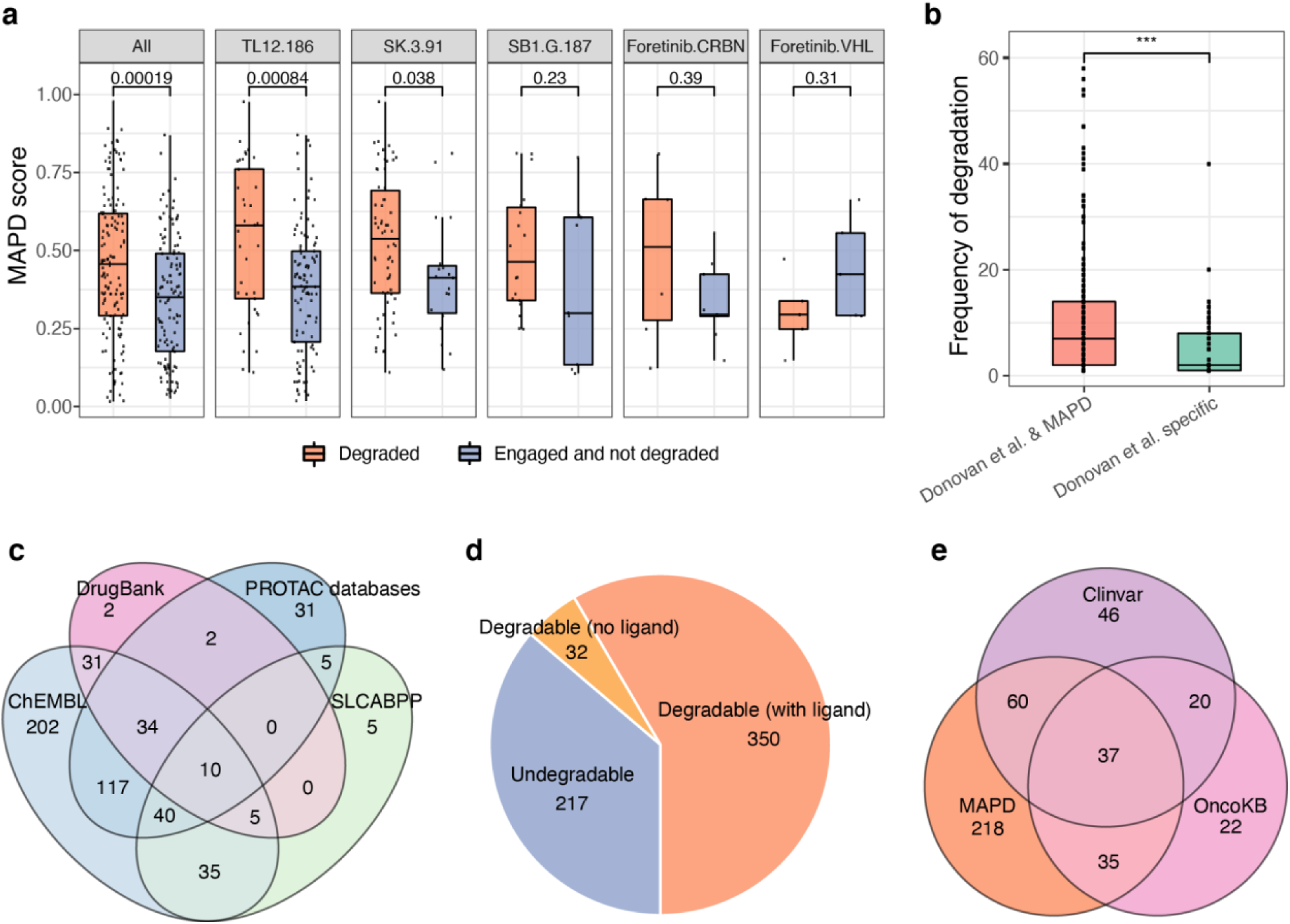
MAPD shows good performance in predicting kinase degradability. Related to Fig. 4. **a**, Box plot showing the MAPD scores of degraded kinases compared to other engaged kinases by each multi-kinase degrader (‘All’ indicates all degraders from Donovan *et al*. study). **b**, Box plot showing the frequency of degradation of degradable kinases identified by both MAPD and Donovan *et al*. and other experimentally degradable kinases (Donovan *et al*. specific). **c**, Venn diagram showing the overlap between ligandable kinases from PROTAC databases (PROTAC-DB and PROTACpedia), DrugBank, ChEMBL, and SLCABPP. **d**, Pie chart showing the number of degradable kinases (with/without ligand) and undegradable kinases from MAPD predictions. **e**, Venn diagram showing the overlap between degradable kinases identified by MAPD, oncogenic kinases reported in the OncoKB, and kinases associated with other human disease reported in the ClinVar database.

**Extended Data Fig. 4.**
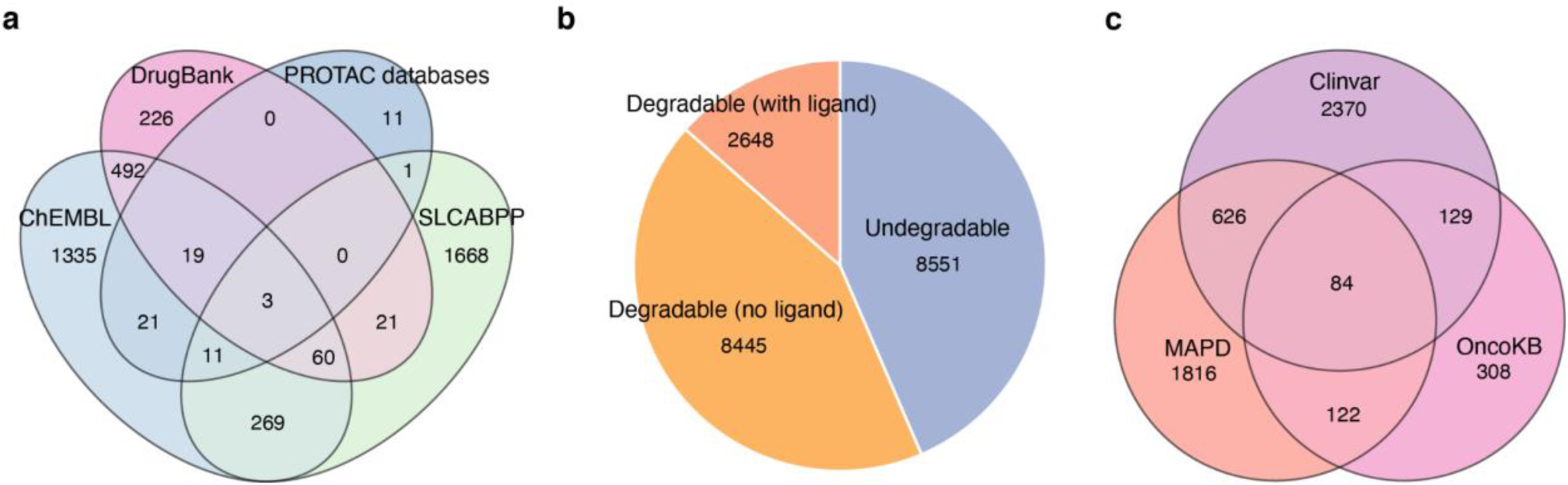
MAPD predicts degradability proteome-wide. Related to Fig. 5. **a**, Venn diagram showing the overlap of ligandable non-kinase proteins from PROTAC databases (PROTAC-DB and PROTACpedia), DrugBank, ChEMBL, and SLCABPP. **b**, Pie chart showing the number of degradable non-kinase proteins (with/without ligand) and undegradable non-kinase proteins from MAPD predictions. **c**, Venn diagram showing the overlap between degradable non-kinase proteins predicted by MAPD and disease-causing proteins reported in the OncoKB and ClinVar database.

**Extended Data Fig. 5.**
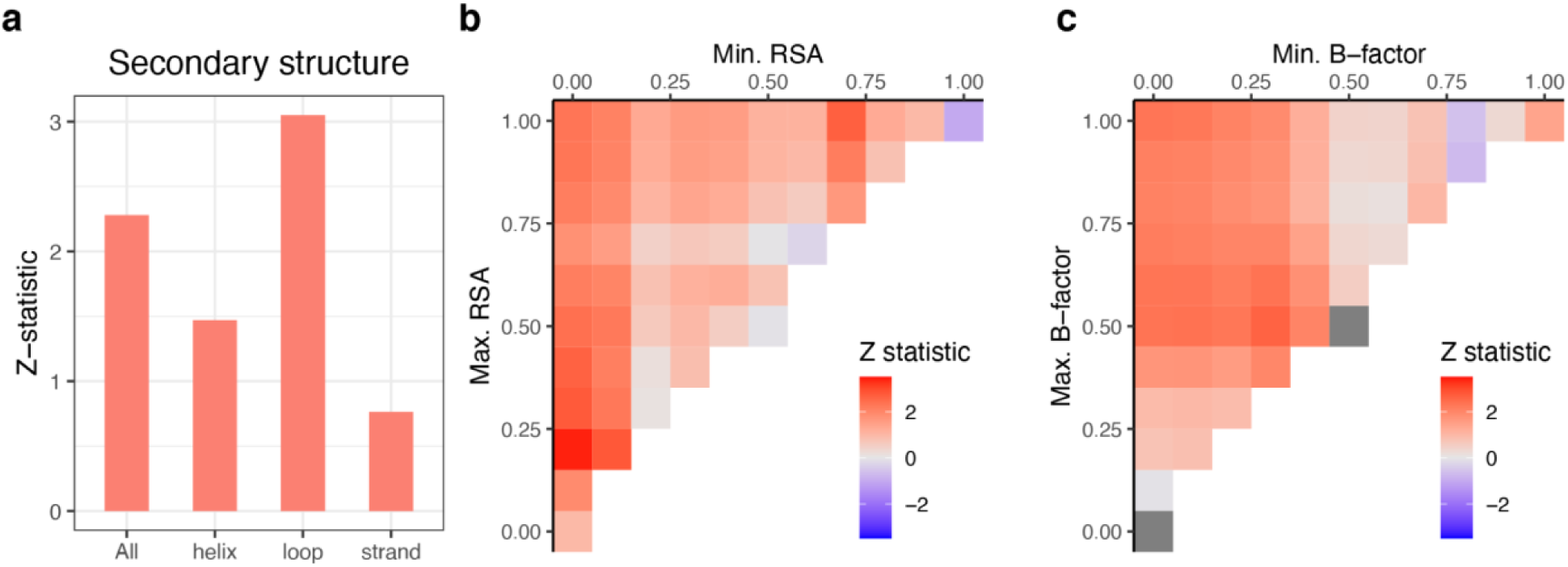
Local structural properties of a Ub site are not informative for predicting protein degradability. **a**, Bar plot showing the Wilcoxon z-statistics that indicate the association between protein degradability and Ub sites in each specific secondary structure. The “All” indicate the total resolved Ub sites in protein structures. **b**, Heatmap showing the Wilcoxon z-statistics that indicate the association between protein degradability and Ub sites in each specific range of relative solvent accessibility (RSA). The x-axis indicates the minimum RSA of each range, and the y-axis indicates the maximum RSA of each range. **c**, Heatmap showing the Wilcoxon z-statistics that indicate the association between protein degradability and Ub sites in each specific range of b-factor (flexibility). The x-axis indicates the minimum b-factor of each range, and the y-axis indicates the maximum b-factor of each range.

**Extended Data Fig. 6.**
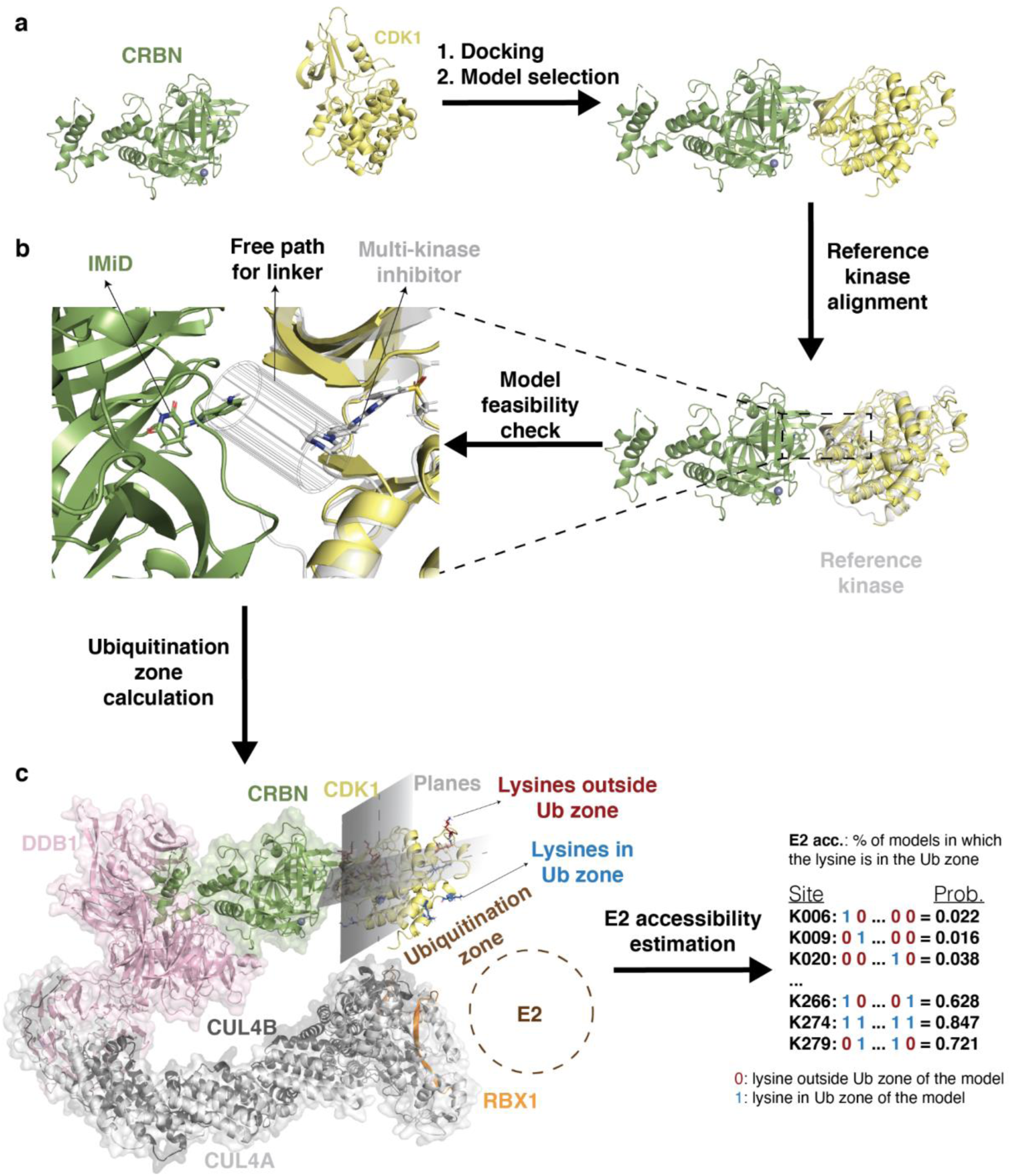
Assessment of E2 accessibility of Ub sites. Related to Fig. 6. **a**, Diagram showing the protein–protein docking process. All kinases were first aligned at their ATP binding pocket to a reference kinase, CDK2 (1AQ1). Next, the aligned kinases were positioned in an arbitrary (but similar) orientation around the ligand-binding pocket of CRBN-Lenalidomide structure (PDB: 5FQD). Here, CDK1 (4Y72) is shown as an example. Local docking was performed, and the 200 top-scoring models were selected for further evaluation. **b**, For every docked model, the feasibility of ternary complex formation with a PROTAC was tested by aligning CDK2 with a multi-kinase inhibitor (TAE) and checking whether a free path for a linker exists. As multiple linkers of different lengths and rigidities were involved, a broad cylinder was used to estimate all linker conformations. **c**, For models where it was feasible to build a ternary complex with a PROTAC, Ub sites in the ubiquitination zone were estimated by drawing two planes with respect to the position of CRBN and the target kinase. The sites lying in the quadrant facing the putative position of the E2, estimated by the placement of RBX1 are considered accessible. For each Ub site, the fraction of feasible models with the site in the ubiquitination zone was used as a probability to measure its E2 accessibility.

**Extended Data Fig. 7.**
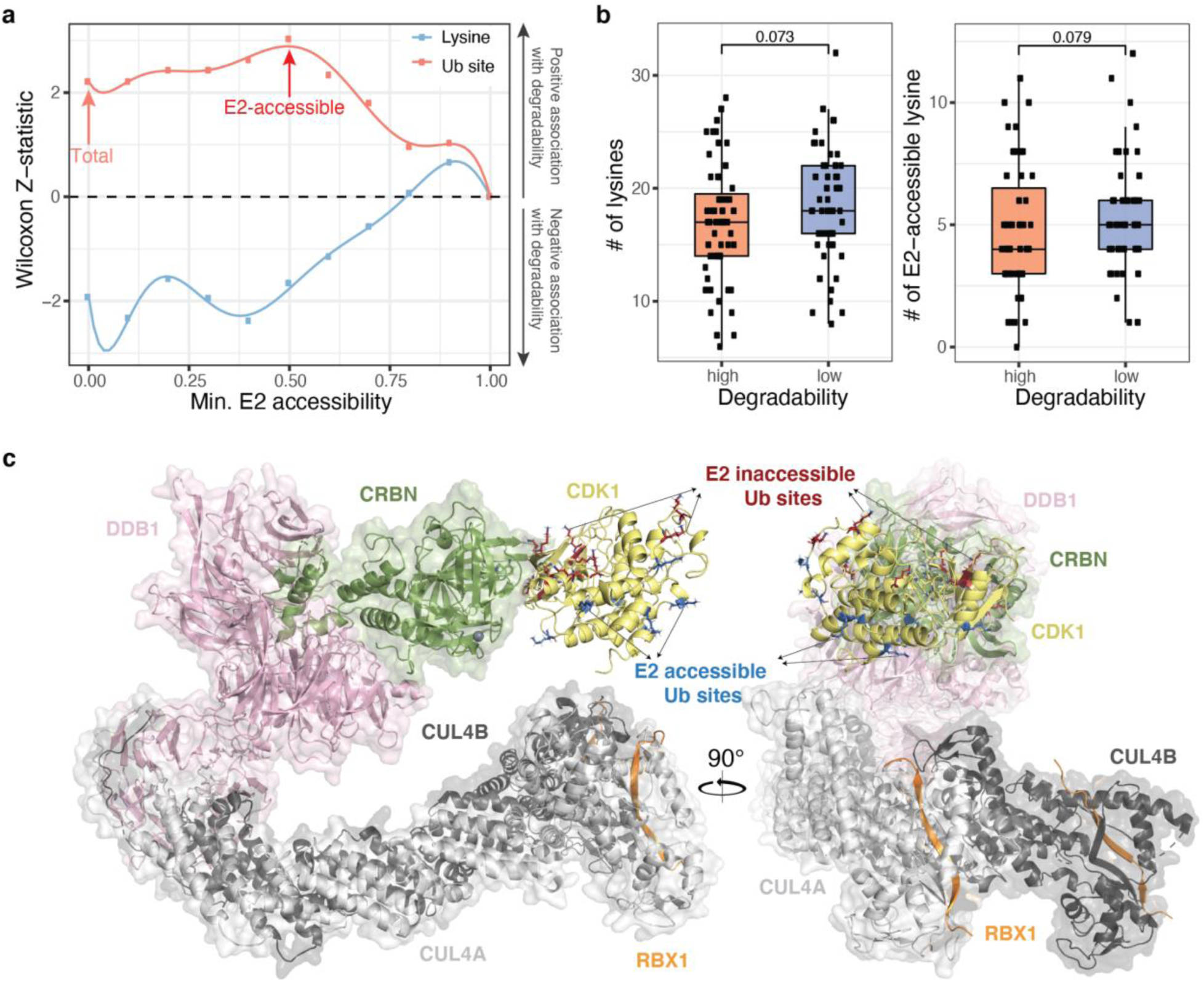
E2-accessibility of Ub sites is associated with protein degradability. Related to Fig. 6. **a**, Smooth line showing the association between protein degradability and the number of E2-accessible Ub sites/lysine residues (E2 accessibility greater than a certain threshold). The x-axis shows the threshold of E2 accessibility for selecting E2-accessible lysine/Ub sites, and the y-axis shows the Wilcoxon z-statistics indicating the association between kinase degradability and the number of lysine/Ub sites with a E2 accessibility greater than a certain threshold. A positive Wilcoxon z-statistic indicates the positive association between protein degradability and the number of lysine/Ub sites, while a negative Wilcoxon z-statistic indicates the negative association between protein degradability and lysine/Ub sites. The salmon arrow points to the association between kinase degradability and the total number of Ub sites, while the red arrow points to the association between kinase degradability and the number of E2-accessible Ub sites (accessible to E2 in more than 50% docking models). **b**, Box plot showing the association of kinase degradability with total number of lysine residues (left) and E2-accessible lysine residues (right) in the kinase targets. The E2-accessible lysine residues (E2 accessibility >=0.5) were defined as the lysine residues lying in the ubiquitination zone of more than 50% feasible models. **c**, Docking model of the ternary complex of CRL4^CRBN^ and the target kinase CDK1. Overlay of CUL4A (PDB: 4A0K) and CUL4B (4A0L) superimposed on DDB1 WD repeat beata-propeller B (4A0K), with CRBN (5FQD) superimposed DDB1 WD repeat beta-propellers A and C demonstrates high flexibility of the CUL4 arm of the E3 ligase. The RBX1 fragment was used to estimate the position of the E2 enzyme and corresponding ubiquitination zone in the target protein CDK1. The model of CDK1 (4Y72) docked to CRBN is shown in yellow, and the predicted E2- accessible and E2-inaccessible Ub sites are highlighted in blue and red, respectively.

## Supplementary Tables

Table 1: A list of protein-intrinsic features.

Table 2: Forward feature selection result for each model.

Table 3: MAPD predictions, ligandability, and disease associations of human proteins.

Table 4: Accessibility of Ub sites to the E2 enzyme in kinase docking models.

## Materials and Methods

### Kinase degradability data

We collected 151 quantitative proteomics data measuring the changes of protein abundance in response to treatment of 85 unique multi-kinase degraders (degraders with allosteric linkers are excluded)^51^. We used the limma package to perform differential protein expression analysis comparing the degrader treated samples with the DMSO treated samples. For each protein, we calculated the frequency of degradation as the number of experiments in which the protein is significantly down-regulated (FC (fold change)>1.25 and p-value<0.01). Furthermore, to aggregate the results of multiple replicates for each degrader, we aggregated log2FC from replicate experiments using Stouffer’s Z-score and corresponding p-values using Fisher’s method. We then counted the number of unique degraders that can degrade each protein (Stouffer’s Z-score< log2(1.5) and Fisher’s p-value<0.01). We collected 5 KiNativ profiling data and 2 KinomeScan data from published studies^51, 52^, which profiled target engagement of five multi-kinase degraders, including TL12-186, SK-3-91, SB1-G-187, DB0646, and WH-10417-099^51, 52^. A KinomeScan score smaller than 15 or a KiNativ score greater than 35 indicate strong drug-target engagement.

### Definition of high-degradability and low-degradability kinases

We defined highly-degradable kinases as those degraded by at least five different multi-kinase degraders (50 kinases), and lowly-degradable kinases that were engaged by at least one multi-kinase degrader, quantified in more than 10% underlying global proteomic experiments, but not degraded (76 kinases). The high-degradable kinases and low-degradable kinases are used throughout the study to investigate the association between protein degradability and protein-intrinsic features.

### Protein-intrinsic features

We built more than 42 protein-intrinsic features spanning post-translational modifications (PTM)^68^, protein stability generated from GPS (global protein stability) profiling^92–94^, protein half-life^95–97^, protein-protein interactions^98, 99^, protein expression, protein detectability^51, 100, 101^, protein length, and others.

#### Post-translational modification (PTM) features

We collected all available post-translational modification (PTM) sites from the PhosphoSitePlus database (02/17/2021)^68^. PhosphoSitePlus includes three types of supports for each PTM site, including LT_LIT (the number of publications supporting the site), MS_LIT (the number of mass spec studies supporting the site), and MS_CST (the number of mass spec studies performed by Cell Signaling Technology supporting the site). We generated two features related to each type of PTM. The first feature (e.g., Ubiquitination_1) refers to the fraction of relevant amino acid residues in a protein (e.g., lysine residues) that have a corresponding reported PTM site (e.g., Ub site), which only needs the support of a single reference for each PTM site (LT_LIT+MS_LIT+MS_CST >0). The second feature (e.g., Ubiquitination_2) is calculated in the same manner, except requires each PTM site to be supported by at least two studies (LT_LIT>1 | MS_LIT>1 | MS_CST >1). We also included the fraction of each likely modified amino acid as additional features, such as LysRatio indicating the fraction of lysine residue in a protein.

#### Protein half-life and protein stability features

We downloaded protein half lives in seven different cell types (B cells, NK cells, Monocytes, Hepatocytes, neurons, Hela, and NIH3T3) from published studies^95–97^. We additionally collected seven global protein stability (GPS) profiling data from three studies^92–94^, which include the stability of full-length proteins in HEK293T cell lines treated with DMSO, MLN4924, dominant negative CRL4, or dominant negative CRL3 and stability of N-terminome and C-terminome peptides of human proteome. All protein half-life data and GPS data were cross-referred for imputing the missing data. The imputation was done by using the impute.knn function (k-nearest neighbor) with default parameters in the impute R package.

#### Protein-protein interaction and protein complex

We downloaded protein-protein interactions (PPI) from the STRING database^98^ and retrieved the high-confidence PPIs using an arbitrary cutoff of experimental score>100 and combined_score>200. The degree of each protein in the PPI network was calculated as an estimation of likelihood of the protein interacting with others. Additionally, curated protein complex annotations were downloaded from the CORUM database^99^ and the number of distinct protein complexes associated with each protein was taken as the estimation of likelihood of a protein being complexed in vivo.

#### Gene and protein expression data

We downloaded RNA-seq data of MOLT4 from the GEO (GSE79253)^102^. RNA expression values were normalized as logarithm Transcripts Per Million (TPM). We retrieved quantitative proteomics data of MOLT4 cell lines from Donovan *et al*., 2020 study^51^. Relative protein abundances were log normalized and centered with a median value of zero per sample. The missing values in the proteomic data were imputed using the impute.knn function (k-nearest neighbor) from the impute R package, with CCLE proteomic data as reference^100^.

#### Protein detectability

We took the frequency of detection of proteins in Donovan *et al*. proteomic datasets as the estimation of protein detectability by mass spectrometry^51^.

#### Other features

We retrieved 20381 reviewed human protein sequences and their length from the UniProtKB database (2021_01). We downloaded Intrinsically disordered regions (IDRs) from the MobiDB database^103^, which includes manually curated annotations and predicted disorder regions. We ranked the IDR annotations based on the four types of evidence, including curated -disorder-priority, derived-missing_residues-th_90, derived-mobile_residues-th_90, and prediction-disorder-mobidb_lite. For each protein, duplicate IDRs were removed for downstream analysis.

### Pairwise correlation of protein-intrinsic features

We computed pairwise spearman correlation of protein-intrinsic features and clustered the features based on the correlation matrix using hierarchical clustering with Euclidean distance measure and complete linkage. The data are visualized using the ComplexHeatmap R package^104^.

### Association between protein degradability and features intrinsic to protein targets

We tested each feature’s difference in 50 high degradability kinases and 76 low degradability kinases using the wilcox.test function in R and computed the Z-statistic using the wilcoxonZ function in the rcompanion R package. We used the same method to test the association between protein degradability and protein-intrinsic features in each kinase family.

### Model-based Analysis of Protein Degradability (MAPD)

We sought to build a classification model to predict protein degradability from intrinsic protein features. We tried six different machine learning models, including linear-kernel SVM (kernlab), radial-kernel SVM (kernlab), random forest (randomForest), K-nearest neighbors, logistic regression (LiblineaR), and naive bayes (naivebayes). For each model, we performed feature selection and then selected the best model trained on a set of best-performing features.

#### Forward feature selection

We performed recursive forward feature selection for six machine learning methods separately. In each iteration, we add a feature which improves the model performance most. The performance is computed as the area under Precision-Recall Curve (auPRC) based on 20-fold cross-validation. This process is stopped when the addition of a new feature does not further improve the performance.

#### Feature importance

We evaluated the importance of features in MAPD using the varImp function in the caret R package^105, 106^, which computes the feature importance on permuted out-of-bag samples based on mean decrease in the accuracy.

#### Performance evaluation

To evaluate the performance of each model involved in the study, we collected prediction scores of all proteins from cross validation and computed the area under the Receiver Operating Characteristic curve (auROC) using the roc function from the pROC package^107^ and Precision-Recall curve (auPRC) using the pr.curve from the PRROC package in R^108^.

#### Single feature evaluation

For each individual feature, we trained a logistic model. For the combination of features, we trained random forest models. Finally, we compared the model performance based on 20-fold cross validation.

#### Final model training for predictions outside of the kinome

We used the caret package for parameter tuning and final model training. We evaluated the model tuning parameters based on leave-one-out cross-validation (method = “LOOCV” in the trainControl function), with the F1 score as performance metric (metric = “F” in the train function, summaryFunction = prSummary in the trainControl function). With the optimal parameters (mtry = 2), we trained a final random forest model including 20,000 trees (ntree = 20,000) with 5 minimum node sizes (nodesize = 5).

### Prediction

We predicted the degradability of all human proteins using the final random forest model. For kinases included in the training, we took the average prediction scores collected from three repeated 20-fold cross-validation. Based on the cross-validation, we chose a cutoff (0.2327) that leads to the highest F1 score. A protein is predicted to be degradable if it has a MAPD score greater than the cutoff. To account for potential biases from missing feature data, we scored the feature completeness for each protein using a weighted sum score with the formula: 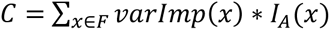. The *F* variable represents the feature set, and *x* represents each feature in the feature set. The function *varImp* (*x*) denotes the scaled feature importance of *x* and the indicator function *I_A_* (*x*) denotes whether *x* is from actual data (1 = actual, 0 = imputed). The *C* represents the feature completeness, with a 0-1 range. A score of 1 indicates all features are from actual data, and a score of 0 indicates all features are imputed.

### Degradable proteins

We collected PROTAC targets with reported degraders in the PROTAC-DB (2021-05-27) and/or the PROTACpedia (2021-07-08)^34, 35^. For evaluation purposes, the targets from Donovan *et al*. study were removed from the PROTAC databases (including PROTAC-DB and PROTACpedia). This resulted in 65 kinases and 65 proteins outside of the kinome. From Donovan *et al*. study, we collected 217 kinases degraded by at least one multi-kinase degrader as ‘degraded’ and all the others detected in the same datasets as ‘not degraded’^51^. We collected 1,336 PROTACtable targets, including the Clinical Precedence targets, Discovery Opportunity targets, and Literature Precedence targets from the PROTACtable genome^74^. We collected 24 IMiD targets from published studies^32^ and assessed their frequency of degradation by 68 CRBN-recruiting multi-kinase degraders from Donovan *et al*. study^51^.

### Protein family

We downloaded the human kinase/kinase-related proteins from four different resources, including KinMap, KinBase, Donovan *et al*. study, and a review article^109–111^. We collected 1,626 human transcriptional factors from a review article^78^.

### Protein ligandability

We downloaded the cysteine reactivity data from the SLCABPP^73^ and assessed protein ligandability using the number of compounds with a competition ratio greater than 4. Besides, we collected protein ligands from the ChEMBL (2021-07-23) and DrugBank database^71, 72^. For any proteins degraded by a multi-kinase degrader or with a ligand recorded in the ChEMBL (2021-07-23), DrugBank, or SLCABPP, we considered it as a ligandable target.

### Protein-disease associations

We considered a protein as a cancer driver if it is an oncogene reported in the OncoKB or it is predicted as an oncogene by 20/20+ algorithm. 20/20+ analysis was performed on the aggregated pan-cancer dataset with default parameters. Genes with an oncogene score greater than 0.5 are considered oncogenes. To annotate potential protein targets associated with other human diseases, we also downloaded the variant-disease association from the ClinVar database^76^ (2021-04-20). For quality control, we removed annotations of likely loss-of-function variants, including indel, deletion, insertion, and microsatellite, as well as some uncertain annotations with key words like ‘conflicting’, ‘protective’, ‘uncertain’, ‘benign’, and ‘not’. This resulted in 3,415 proteins associated with human diseases reported in the ClinVar database.

### Structural properties of lysine residues and Ub sites

We downloaded protein structures of human models or homology models from PDB^112^, SWISS-MODEL^113^, and ModPipe^114^. The detailed data cleaning and processing have been described in Tokheim *et al*. study^115^. Protein structures were analyzed using the DSSP program^89^ in bio3d R package^116^, which returns the solvent accessibility and secondary structure of each residue.

### Protein-protein docking

We downloaded protein structures of 323 kinases from the PDB. In cases where multiple structures were available, the largest structure was chosen. They were aligned to CDK2 (PDB: 1AQ1)^117^, a reference kinase, to ensure that the kinase domain was present. 251 kinase structures were alignable with root-mean-square deviation less than 3.5 Å near the ATP-binding pocket. Next, the aligned kinases were positioned in an arbitrary (but similar) orientation around the ligand-binding pocket of CRBN–Lenalidomide structure (PDB: 5FQD)^14^. Using Rosetta v.3.12^118^ and RosettaDock v.4.0^119^, we performed 5,000 independent local docking with different starting points and perturbation of 3 Å and 8° (all options listed below). Models were evaluated by the interface score metric (I_sc) and the 200 lowest-scoring models were selected for further evaluation.

### E2 accessibility of lysine residues

We assessed the accessibility of solvent-exposed lysine residues to the E2 enzyme by calculating the fraction of protein-protein docking models among the 200 lowest-scoring models that could fit a PROTAC and in which the lysine residues are in the ubiquitination zone of the E2 enzyme. All lysines with any atom having >2.5 Å^2^ exposed surface area were considered solvent exposed. The ability of the ternary complex to fit a PROTAC was assessed by aligning CDK2 with CDK4 inhibitor (PDB: 1GIJ)^120^ to the kinase and calculating if there was a free path available between the N3 atom Lenalidomide and C26 atom of the CDK4 inhibitor to build a linker. If a cylinder of radius 1 Å and length <14 Å could be constructed between the aforementioned atoms with less than 2 protein backbone or compound atoms (except neighboring atoms) inside the cylinder, we estimated that there exists a free path to build a linker, and hence fit the PROTAC. To assess which lysine residue lie within the ubiquitination zone of the E2, we constructed two planes to split up space into quadrants. The ‘vertical’ plane passes through half the distance between the CRBN edge facing the kinase and the center-of-mass of the kinase. The ‘horizontal’ plane is approximately perpendicular to the vertical plane and passes through the center-of-mass of the kinase. The lysine residues lying in the quadrant facing the putative position of the E2 are considered accessible. Finally, if the lysine residue was more than 60 Å away from the Lenalidomide or the C_α_–C_β_ vector points in the direction opposite of the putative E2 site, the residue was considered inaccessible.

### Association between protein degradability and characteristics of Ub sites

We first counted each protein’s lysine residues/Ub sites in different secondary structures (coil, strand, and loop), and then tested whether there is a difference between highly-degradable and lowly-degradable kinases using the Wilcoxon z-statistics. Similarly, we assessed the associations between kinase degradability and the number of lysine residues/Ub sites with a specific range of solvent accessibility or B-factor. A positive Wilcoxon z-statistic indicates the positive correlation between kinase degradability and the number of Ub sites/lysine residues in the proteins.

We also tested the association between kinase degradability and the number of E2-accessible Ub sites/lysine residues (E2 accessibility greater than a specific threshold) in each protein. To further demonstrate the specific importance of E2-accessible Ub sites, we randomly shuffled the Ub sites among all lysine residues and re-evaluated the association between kinase degradability and the number of E2-accessible Ub sites in each kinase. We generated a null distribution by repeating the shuffling process for 10,000 times and calculated the p-value by counting the percentage of shuffling that led to a higher Wilcoxon z-statistic than the observed Wilcoxon z-statistic.

### Data and software availability

The R package is stored on github: https://github.com/liulab-dfci/MAPD. The source code for reproducible data analysis is stored on github: https://github.com/liulab-dfci/Degradability2021. All relevant data and results are accessible at http://mapd.cistrome.org.

